# Oxytocin via oxytocin receptor excites neurons in the endopiriform nucleus of juvenile mice

**DOI:** 10.1101/2022.03.04.483043

**Authors:** Lindsey M. Biggs, Elizabeth A.D. Hammock

## Abstract

The neuropeptide oxytocin (OXT) modulates social behaviors across species and may play a developmental role for these behaviors and their mediating neural pathways. Despite having high, stable levels of OXT receptor (OXTR) ligand binding from birth, endopiriform nucleus (EPN) remains understudied. EPN integrates olfactory and gustatory input and has bilateral connections with several limbic areas. Because the role of OXTR signaling in EPN is unknown, we sought to provide anatomical and electrophysiological information about OXTR signaling in mouse EPN neurons. Using in situ hybridization, we found that most EPN neurons co-express *Oxtr* mRNA and the marker for VGLUT1 and are thus glutamatergic cells. Based on high levels of OXTR ligand binding in EPN, we hypothesized that oxytocin application would modulate activity in these cells as measured by whole-cell patch-clamp electrophysiology. Bath application of OXT and an OXTR specific ligand (TGOT) increased the excitability of EPN neurons in wild-type, but not in OXTR-knockout tissue. These results show an effect of OXT on a presumably glutamatergic cell population within EPN. Given the robust, relatively stable OXTR expression in EPN throughout life, OXTR in this multi-sensory and limbic integration area may be important for modulating activity in response to an array of social or other salient stimuli throughout the lifespan and warrants further study.

**Significance statement:** There is a high level of oxytocin receptor (OXTR) expression in the mouse endopiriform nucleus (EPN) throughout development, however little is known about the effect of oxytocin on these neurons. We show here that *Oxtr* mRNA co-expresses mainly with VGLUT1, a glutamatergic cell marker. Using OXTR-EGFP mice to identify EPN, we show that OXT has a mainly excitatory effect on EPN neurons. Thus, activity in EPN neurons may be modulated by OXT during exposure to salient or social stimuli throughout development and this could affect development of behavioral responses during social exposure.

## Introduction

Oxytocin (OXT) is a nine amino acid neuropeptide that acts through the oxytocin receptor (OXTR) to generate physiological responses in adult mammals, such as facilitating milk let down for lactation, and labor and delivery. OXT also acts in the central nervous system to modulate social behavior by facilitating social learning and memory ^1,2^, social recognition ^3–5^, social reward ^6,7^, pair-bond formation ^8,9^, parental care ^10–13^, sexual behavior ^14–17^, and by reducing aggression ^18,19^. Specifically, activation of OXT producing neurons in the paraventricular nucleus (PVN) of the hypothalamus and subsequent OXT release increases social behavior, while reduced activation of these neurons results in a decrease in social behavior ^20^. The effect of OXT on many areas of the central nervous system has been investigated, including the hippocampus where it modulates learning and memory ^21–24^, areas that receive sensory input (i.e. olfactory bulb, anterior olfactory nucleus, primary auditory cortex) where it modulates sensory input based on the social salience of the signal ^5,11,25^ and amygdalar areas in order to decrease fear-related behavior ^26^ and facilitate social recognition ^2,27,28^.

One brain area that has received little attention that may play a role in the OXT modulation of behavior is the endopiriform nucleus (EPN). Previous autoradiographic binding and immunohistochemical studies have shown that, unlike some other areas of the developing mouse brain that display transient oxytocin receptor (OXTR) expression ^12,29,30^, OXTR expression in the EPN is dense and remains relatively stable throughout development from postnatal day 0 (P0) to adulthood (reviewed in Figure 1) ^29,31^. Despite robust OXTR expression throughout development, the effect of OXT on the activity of EPN neurons has not been investigated.

**Figure 1.**
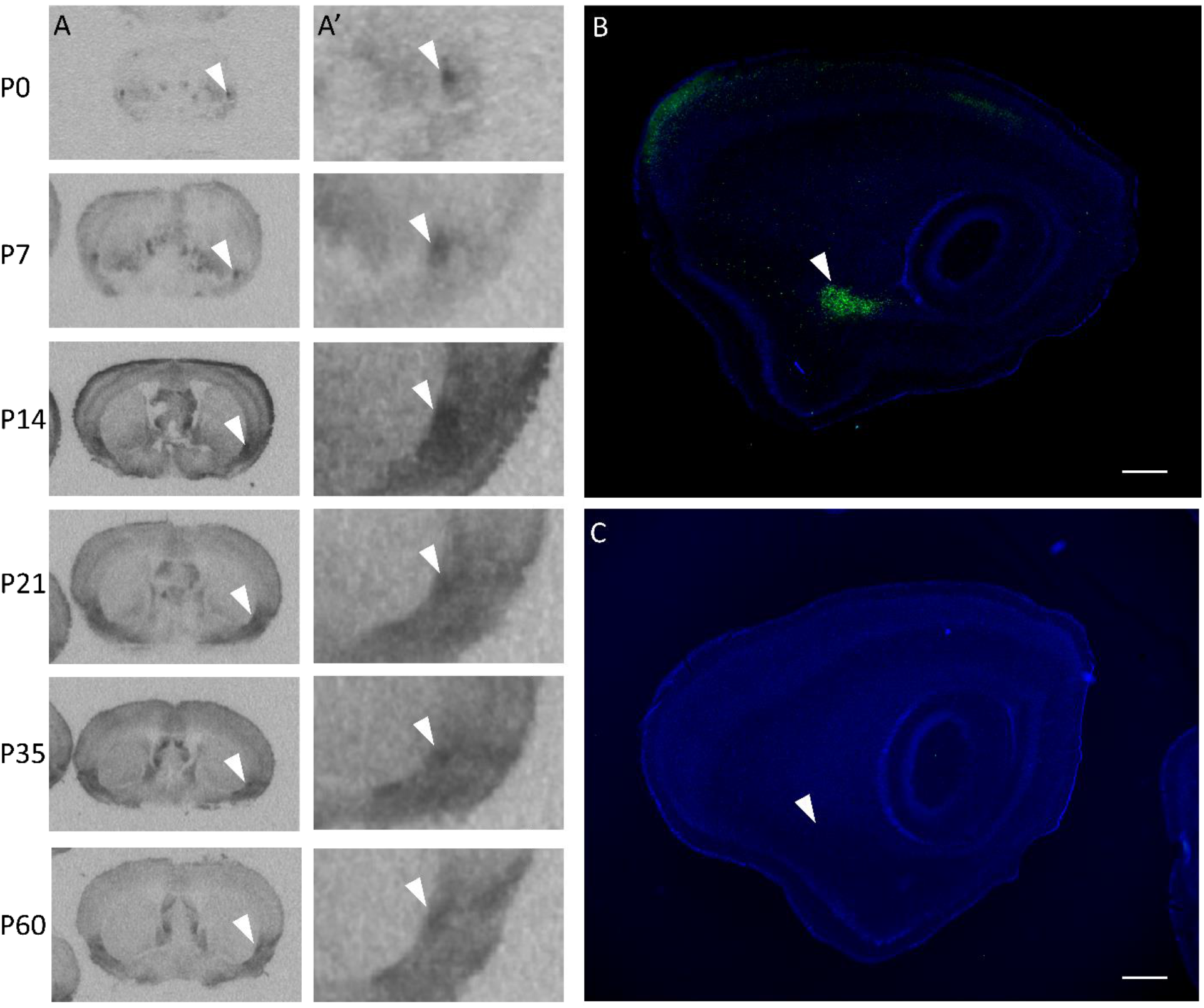
OXTR is produced by and present in the endopiriform nucleus (EPN) across development in mice. A) Previously unpublished autoradiographic images selected from prior work by Hammock and Levitt (2013) demonstrate OXTR ligand binding in the EPN from birth in C57BL/6J mice. White arrowheads in A and A’ (4x zoom) point to the dorsal EPN in the coronal plane at post-natal day 0 (P0), P7, P14, P21, P35, and P60. OXTR ligand binding was present in both males and females at all ages examined. B) Sagittal view of Oxtr-EGFP reporter mouse at P21 (enhanced with immunofluorescence for EGFP, counterstained with DAPI) demonstrates the expression of the EGFP reporter throughout the rostro-caudal extent of the EPN. C) There is no EGFP signal in a transgene negative sample. White arrowheads in B and C show the EPN at P21. Scale bar = 0.5 mm

To investigate a role for OXT signaling at OXTR in the mouse EPN, we used anatomical methods to identify the cell-type that predominately expresses OXTR in EPN. Whole-cell electrophysiology was used to investigate the effect of OXT on EPN neurons in male and female mice at P21-P28. We used an OXTR specific ligand to validate the OXTR-dependent effect and confirmed the absence of an effect in OXTR knock-out tissue.

## Methods

### Animals

All protocols and procedures were performed after approval from the Animal Care and Use Committee at Florida State University in accordance with state and federal guidelines and reported in accordance with ARRIVE guidelines. All mice were bred in the animal facilities at Florida State University. Mice expressing enhanced green fluorescent protein (EGFP+) upstream of the oxytocin receptor gene (OXTR-EGFP) in a mixed FVB/N x Swiss-Webster background were originally obtained from GENSAT (GO185Gsat/Mmucd) ^32^. Mice carrying the *Oxtr* allele (*Oxtr*^*tm*1.1*Knis*^) ^33^ were fully backcrossed to C57Bl6j. For OXTR-EGFP experiments, a male OXTR:EGFP transgene positive mouse was paired with an EGFP negative female. To generate *Oxtr* ^-/-^ mice in the context of the OXTR-EGFP reporter, the OXTR-EGFP line was crossed with the *Oxtr* line to generate EGFP+/*Oxtr*^*+/-*^. These EGFP+/*Oxtr*^*+/-*^ mice were then bred with *Oxtr*^*+/-*^ to generate offspring that could be EGFP+ and also *Oxtr*^+/+, +/-^, or ^-/-^. Breeder cages were checked daily for litters and the first appearance of a litter was indicated as post-natal day 0 (P0). All litters were weaned into male and female specific cages at P21. All animals were housed in the vivarium under a 12/12 light cycle and provided ad libitum food and water.

### Receptor autoradiography

Autoradiographic images of OXTR ligand binding were generated in a previously reported study ^29^. Previously unpublished images were selected from that dataset to specifically illustrate the presence of OXTR in the EPN across development.

### EGFP immunohistochemistry

To better visualize the EGFP reporter to examine reporter expression in the EPN, 40 μm free floating sagittal paraformaldehyde fixed brain sections from transcardially-perfused P21 mice were first washed four times 15 minutes in PBS, one time 20 minutes in 0.5% sodium borohydride in PBS, 4 times 5 minutes in PBS, then incubated overnight at room temperature with Chicken anti-EGFP (1:5,000; Abcam ab13970, lot# GR3190550-18) in 1% goat serum and 0.3% Triton X100 in PBS. After 4 times 15 minute washes in PBS, tissues were incubated for 2 hours at room temperature in Alexafluor-488 conjugated donkey anti-chicken (1:500; Jackson ImmunoResearch 703-546-155, lot# 142461) in 1% goat serum and 0.3% Triton X100 in PBS. Sections were washed five times 5 minutes in PBS, incubated in 0.1 μg/mL DAPI for 5 minutes, washed five times 5 minutes in PBS then mounted onto gelatin subbed slides, coverslipped with Vectashield (Vectorlabs H-1000) and visualized with a Keyence fluorescence microscope. EGFP transgene positive tissue was compared to EGFP transgene negative tissue.

### In situ hybridization

RNAscope in situ hybridization was performed using the RNAscope 2.5 HD kit (Advanced Cell Diagnostics, Inc). The following probes were used: *Oxtr* mRNA:Oxtr-E4_C2, #411101-C2 (targeting sequence 1198-2221 of NM_001081147.1, *GenBank*); *Gad1*:#400951 (targeting sequence 62-3113 of NM_008077.4, *GenBank*); *Gad2*: #439371 (targeting sequence 552-1506 of NM_008078.2, *GenBank*); *SLC17a7* (VGLUT1): #416631 (targeting sequence 464-1415 of NM_182993.2, *GenBank*); *SLC17a6* (VGLUT2): #319171 (targeting sequences 1986-2998 of NM_080853.3, *GenBank*). Tissue was collected from a female C57BL/6J mouse (P28), sectioned at 20 μm and stored at -80°C until processing.

### Biocytin immunostaining

During initial pilot recordings, biocytin (Biotium 90055, 1 mg/ml) was added to the intracellular fluid to verify recording location. After recording, tissue was immediately placed into 0.4% paraformaldehyde for approximately 24 hours at 4°C, then transferred to phosphate buffered saline (PBS, pH 7.4) until immunostaining. After several PBS washes, tissue sections were incubated overnight in PBS-triton, 3% bovine serum and HiLyteFluor Texas Red-conjugated streptavidin (AnaSpec, Inc. 60671, Lot# 57051-066-078; 1:1,000). After several washes in PBS-triton, slices were incubated in DAPI (1:1000) for 10 minutes, mounted and coverslipped with Vectashield (H-1000).

### Acute slice preparation

Male and female mice (P21-P28) were used for all electrophysiology experiments. After decapitation, brains were quickly removed from the skull of un-anaesthetized mice and placed into oxygenated (95% O_2_, 5% CO_2_), ice cold sucrose solution containing (in mM): NaCl 83; NaHCO_3_ 26.2; NaH_2_PO_4_ 1; MgCl_2_ 33; CaCl_2_ 0.5; glucose 22; sucrose 72. EGFP expression was verified at the time of dissection by examining the trigeminal nuclei in the base of the skull with a fluorescent light as these ganglia show high expression of OXTR-EGFP fluorescence ^34^. A block of tissue containing the EPN was dissected out and placed into a vibratome containing the sucrose solution, hemisectioned, and coronal slices (300 μm) containing the EPN were sectioned. Slices were then transferred to a holding chamber and incubated at 33°C for 30-45 minutes in oxygenated (95% O_2_, 5% CO_2_) artificial cerebrospinal fluid (aCSF) containing (in mM): NaCl 119; NaHCO_3_ 26.2; KCl 2.5; CaCl_2_ 2.5; NaH_2_PO_4_ 1; MgCl_2_ 1.3; glucose 22. Slices were then allowed to equilibrate to room temperature in the same holding chamber for at least 30 minutes, then individual sections were transferred to the recording chamber mounted on an Olympus BX51WI upright microscope equipped with brightfield and fluorescent capabilities.

### Whole cell electrophysiology

Whole cell current-clamp recordings were obtained from neurons located within the OXTR-EGFP+ EPN. The EPN was identified by a dense expression of EGFP identified via excitation with blue light in the OXTR-EGFP mouse line and by using neuroanatomical landmarks described below. Under high magnification on our recording apparatus, there were OXTR-EGFP+ cells and similarly sized cells that did not have EGFP levels detectable by our camera. Healthy cells from both groups were patched under brightfield illumination and showed no significant differences in electrophysiological responses to the experimental conditions so responses were pooled together for these analyses. For whole-cell recordings, glass pipettes (3-8 MΩ, 1B150F-3, World Precision Instruments) were pulled using a horizontal puller (Sutter Instruments, Inc) and had an initial resistance of 3-7 MΩ.

Pipettes were filled with an intracellular fluid that contained (in mM): KMeSO_3_ 145; MgCl_2_-6 H2O 1; Hepes 10, EGTA 1.1, Na-ATP 2; Na-GTP 0.4 (pH 7.3-7.4, ∼280 mOsm). Whole-cell configuration was performed only after acquiring a GΩ seal and cells with an initial access resistance higher than 25 MΩ were excluded from analysis. All recordings were performed with a Multiclamp 700B amplifier (Molecular Devices) and controlled by PClamp software (Version 10, Molecular Devices). Capacitive transients were automatically compensated and all data was acquired at 10 kHz. An experimentally defined junction potential of -11 mV was applied to all traces.

All agonists and antagonists were bath applied prior to, during and after addition of oxytocin or the OXTR specific ligand, threonine^4^ glycine^7^ oxytocin (TGOT), in order to obtain accurate baseline, experimental and washout data respectively. The following antagonists were used: GABA receptor antagonist, picrotoxin (100 μM, P1675, Sigma-Aldrich); NMDA glutamate receptor antagonist, APV (10 μM, K3375, Sigma-Aldrich); AMPA/Kainate glutamate receptor antagonist, DNQX (5 μM, #189, Tocris Bioscience); sodium channel blocker, Tetrodotoxin (TTX, 1 μM, Tocris Bioscience, #1078). The oxytocin neuropeptide (200 nM, 06379, Sigma-Aldrich) and the oxytocin-receptor specific agonist, TGOT (200 nM,Bachem, H-7710) were also bath-applied as indicated in the Results section. To avoid any confounding effect of prior exposure to agonist/antagonist bath solutions, only one cell was recorded per slice. Once the tissue was exposed to an experimental bath solution, the tissue was discarded after recordings.

### Data Quantification

All quantification was performed in Clampfit 10.7. Resting membrane potential was calculated as the average membrane potential over 10 seconds prior to current injection of every sweep collected during the experiment. Resistance was calculated from the steady state voltage response to -50 pA current injection. Number of evoked action potentials was in response to 50-150 pA current injection determined for each cell to elicit 3-10 action potentials per sweep under baseline conditions.

Statistical analyses were performed using Graphpad Prism 8. All electrophysiological data was analyzed as a one-way repeated measures ANOVA with Greenhouse-Geisser correction followed by Dunnett multiple comparisons post-hoc test comparing drug application and washout to baseline conditions. Graphical analysis of the residuals was used to test the statistical assumptions for the ANOVA. Cohen’s d was calculated by dividing the mean difference between the two group by the pooled standard deviation to determine the effect size. All data are presented as mean +/- SEM. The sample size for statistical analysis is at the level of the individual cell/slice and is indicated along with the number of animals used in the Results section and Supplementary Table 1. Supplementary Table 1 also contains the results for each statistical assay. The data used for each statistical test is included in Supplementary tables 2-14.

## Results

Previously published autoradiography experiments have identified areas with high levels of OXTR ligand binding using the highly selective ligand ([125I]-OVTA) in mice. Unpublished images from this published dataset ^29^ demonstrate OXTR ligand binding in the EPN of C57BL/6J mice from post-natal day 0 (P0) through P60 (Figure 1A). To attempt to identify potential sources of the high levels of OXTR ligand binding in the EPN, we performed EGFP immunohistochemistry in perfused tissue from the OXTR-EGFP reporter line. Robust EGFP immunoreactivity was evident within the EPN of P21 sagittal sections of OXTR-EGFP reporter mice (Figure 1B), but not in mice without the EGFP transgene (Figure 1C). Investigation of the cell types that express OXTR was performed using RNAscope in situ hybridization staining to identify co-expression of *Oxtr* and several markers for glutamatergic and GABAergic cell types (Figure 2). Within the EPN, most *Oxtr* mRNA expressing neurons co-express *Slc17a7*, a marker for VGLUT1 (Figure 2C), a glutamate transporter protein located on the membrane of glutamate-containing vesicles ^35^ that is commonly used to identify glutamatergic neurons and is commonly found in the neocortex, hippocampus and basolateral amygdala ^36^. A low level of *Slc17a6* mRNA, a marker for VGLUT2 cells, was also found in the EPN, however, co-localization of *Oxtr* mRNA expressing cells was much lower than that seen with VGLUT1 (Figure 2D). Subpopulations of GABAergic cells can be identified by the presence of two different glutamate decarboxylases, GAD65 and GAD67, enzymes that are responsible for the transition from glutamate to GABA and CO_2_ ^37^. The GAD65 and GAD67 proteins are encoded by two distinct genes, *Gad1* and *Gad2* respectively. Both *Gad1* and *Gad2* mRNA were found in EPN neurons. Although the density of expression for *Gad1* and *Gad2* was lower than that seen for VGLUT1, there was still some co-expression of *Oxtr*-mRNA and both *Gad1* (Figure 2A) and *Gad2* (Figure 2B). Overall, the results of the mRNA expression show that *Oxtr* mRNA is densely expressed in the EPN, such that *Oxtr* mRNA was expressed in almost all cells identified by the hematoxylin counterstain. Further, the majority of these *Oxtr* mRNA expressing neurons are glutamatergic with a small subpopulation of GABAergic *Oxtr*-mRNA neurons within the EPN.

**Figure 2.**
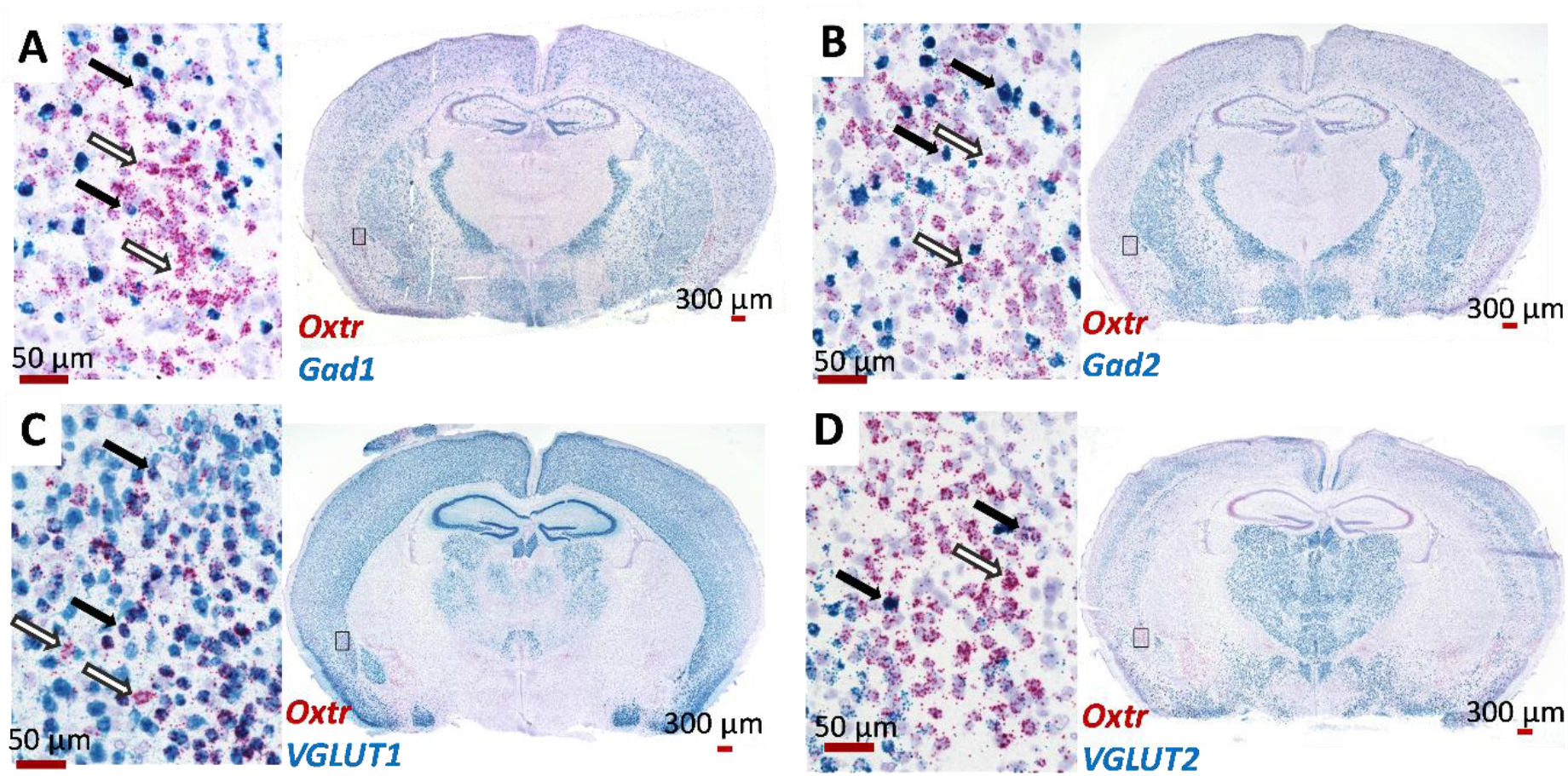
Endopiriform nucleus (EPN) cells that express *Oxtr* mRNA are glutamatergic, with a small population of GABAergic *Oxtr*-mRNA cells. Some neurons that express *Oxtr* mRNA (red) co-express mRNA for the GABAergic markers (black arrows), *Gad1* (A, blue) or *Gad2* (B, blue). However, most *Oxtr*-mRNA expressing cells were negative for these markers (white arrows). Most of the *Oxtr* mRNA expressing neurons in EPN co-express the markers for VGLUT1 (*Slc17a7*) (C, blue, black arrows) with only a few that expressed only *Oxtr* mRNA (C, white arrows). Expression of the marker for VGLUT2 (*Slc17a6*) (D, blue) was low in EPN but some co-expression with *Oxtr* was detected (black arrows).

To further study the EPN neurons, a transgenic mouse line (OXTR-EGFP) was used to easily identify the OXTR-expressing neurons in live tissue slices. This mouse line expresses a green fluorescent protein under the OXTR-promotor region as seen in Figure 1B, 3B and 3C. The EPN is located just lateral to the external capsule and medial to the piriform cortex so these cytoarchitectural landmarks were used to identify the EPN in live tissue slices. The EPN was distinguished from the claustrum by the location of the rhinal fissure with EPN located ventral and claustrum dorsal. During pilot electrophysiological recordings, the location of the patch clamped cell was identified by adding biocytin (1 mg/ml) to the intracellular solution, followed by post-hoc immunostaining. In all pilot recordings, the biocytin filled cell (Figure 3A,C) was located within the EPN as identified by the cytoarchitectural landmarks described above and by OXTR-EGFP expression (Figure 3B, C). All subsequent electrophysiological recordings were performed using the same parameters, however biocytin was not added to the intracellular solutions to limit any potential effect of biocytin on the health or behavior of the cells during the recordings. We then performed whole-cell patch clamp electrophysiology to investigate the effect of oxytocin and an oxytocin receptor selective agonist, TGOT (threonine^4^ glycine^7^ oxytocin) on EPN neurons ^38^. For these experiments, all agonists and antagonists were bath applied at a rate similar to the aCSF drip rate (∼1 drip/sec). In pilot experiments using tetrodotoxin (TTX), evoked action potential firing was monitored after the start of the TTX bath to estimate the time necessary for sufficient fill and turnover of the fluid in the recording chamber. In these pilot experiments, action potential firing was abolished approximately 3-4 minutes after starting the bath application, so all experimental data was collected at 4 minutes and 6 minutes after the start of bath application. In EPN neurons held just above threshold by injecting positive current in current clamp recordings, there was a significant main effect of bath application of oxytocin (200 nM) (One-way repeated measures (RM) ANOVA F (2.112, 10.56) = 6.315, p=0.015, n=6 cells, 3 animals-1 male/2 female). Post hoc comparisons using the Dunnett’s multiple comparisons test (MCT) showed that EPN neurons exhibited more spontaneous action potentials at 4 minutes into OXT bath application compared to baseline levels (MΔ=32.83, SE 6.779, Q=4.843, df=5, p=0.011, d=1.87, Figure 4A, Supplementary Table 1 Line A). The number of spontaneous action potentials fired at 6 minutes was not significantly different from baseline and the cells returned to baseline levels of spontaneous firing after 20 minutes of washout with aCSF (MΔ=-6.667, SE=8.265, Q=0.807, df=5, p=0.768).

**Figure 3.**
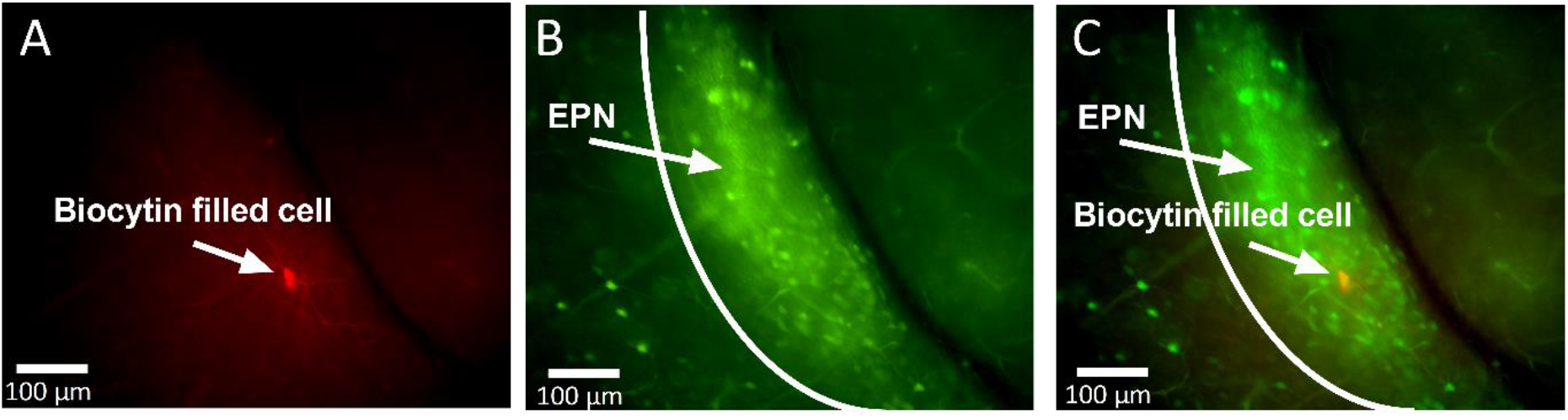
OXTR-EGFP expressing neurons in the endopiriform nucleus (EPN). During preliminary electrophysiological recordings, biocytin was added to the intracellular solution and injected into the recorded cell. Post-labelling of these cells (A) revealed anatomical location of neurons recorded from during electrophysiological experiments. Endogenous expression in the OXTR-EGFP mouse tissue slice (350 μm) shows dense OXTR-EGFP expression in the EPN (B). The combined image (C) shows that the location of the cell recorded from was the OXTR-EGFP rich EPN.

**Figure 4.**
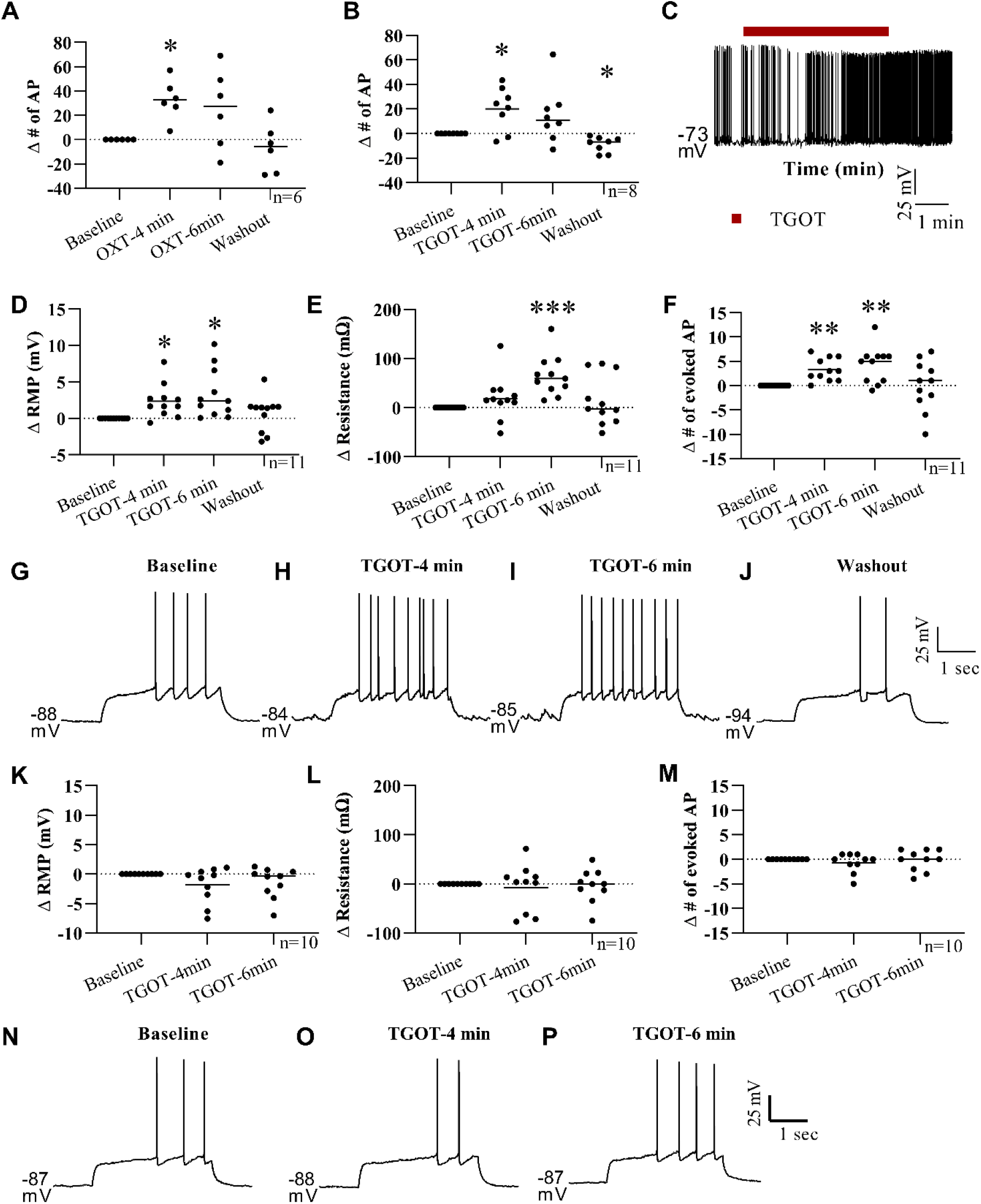
Oxytocin and TGOT have a depolarizing effect on EPN neurons. When held near the action potential threshold in current clamp recordings, bath application of OXT (A, 200 nM) and TGOT (B, 200 nM) elicited a significant increase in the number of spontaneous action potentials at 4 minutes after the start of application. Representative trace (C) shows increase in the spontaneous action potential firing after start of TGOT (red line) bath application which persists for several minutes. In a separate group of cells at their resting membrane potential, the membrane potential (D) was significantly depolarized with an increase in the number of evoked action potentials (F) at 4 and 6 minutes after the start of TGOT bath application. A significant increase in membrane resistance was detected only at the 6 minute timepoint in this experiment (E). Representative traces from one neuron show the increase in evoked action potential firing at 4 and 6 minutes with a return to baseline levels during washout (G-J). In a separate experiment using OXTR-KO EGFP mice, TGOT bath application had no significant effect on the cells membrane potential, resistance or number of evoked action potentials (K-M, representative traces in N-P). * p<0.05; ** p<0.005; *** p<0.001.

OXT not only binds to the OXTR which we have shown is present in the EPN, but it also binds to vasopressin receptors ^39–43^ so it is possible that the effect seen above was due to OXT binding at vasopressin receptors. To avoid binding at vasopressin receptors, the OXTR selective agonist TGOT was used for the remainder of the experiments ^38^. Using the same experimental design as described above, an aCSF solution containing TGOT (200 nM) was bath applied to the tissue slice during whole-cell patch clamp electrophysiological recording of an EPN neuron held just above threshold in current clamp and the number of spontaneous action potentials was quantified before, during and after TGOT application. One-way RM ANOVA indicated a significant main effect of treatment (F (1.483, 10.38) = 8.918 p=0.008, n=8, 2 animals-1 male/1 female). When compared to baseline using Dunnett MCT, EPN cells fired significantly more action potentials at 4 minutes into the bath application of TGOT (MΔ=20.13, SE=6.26, Q= 3.215, df=7, p=0.036, d=1.25) and exhibited significantly fewer spontaneous action potentials than baseline after 20 minutes of aCSF washout (MΔ=-8.75, SE=2.22, Q=3.94, df=7, p=0.014, d=1.35. Figure 4 B,C, Supplementary Table 1 Line B).

To further characterize the effect of TGOT on EPN neurons at the resting membrane potential (RMP), a separate experiment was performed at the cells RMP with a three second positive current injection experimentally determined for each cell such that the current evoked between 2 and 10 action potentials during this time (n=11, 4 animals-2 male/2 female). One-way RM ANOVA indicated a significant effect of TGOT bath application on the EPN cells RMP (F (2.486, 24.86) = 6.165, P=0.006) with a significantly depolarized RMP at both 4 (Dunnett MCT, MΔ= 2.35 mV, SE=0.699, Q=3.362, df=10, p=0.019, d=0.275) and 6 minutes (Dunnett MCT, MΔ=3.355 mV, SE=1.026, Q=3.271, df= 10, p=0.022, d=0.379) after the start of the TGOT bath application compared to baseline, with a return to baseline RMP after the 20 minute aCSF washout (i.e. no significant difference between baseline and washout, MΔ=0.645, SE=0.737, Q=0.875, df=10, p=0.72) (Figure 4D, Supplementary Table 1 Line C). TGOT bath application also had a significant main effect on EPN cells membrane resistance (F (1.880, 18.80) = 5.533, p=0.014) with a significant increase in the resistance at 6 minutes relative to baseline (Dunnett MCT, MΔ=64.93 mΩ, SE=12.35, Q=5.259, df=10, p=0.001, d=0.518) with a return to baseline levels upon washout (MΔ=14.18 mΩ, SE=15.16, Q=0.935, df=10, p=0.682) (Figure 4E, Supplementary Table 1 Line D). A significant main effect of TGOT on the number of experimentally evoked action potentials was also found (F (1.713, 17.13) = 5.426, p=0.018) with significantly more evoked action potentials at both 4 (Dunnett MCT, MΔ=3.273, SE=0.715, Q=4.579, df=10, p=0.003, d=1.185) and 6 minutes (MΔ=4.273, SE=1.129, Q=3.785, df=10, p=0.009, d=1.739) after the start of the TGOT application with no significant difference between washout and baseline levels (MΔ=0, SE=1.543, Q=0, df=10, p>0.999) (Figure 4F-J, Supplementary Table 1 Line E).

To further confirm that OXTR is necessary for the depolarizing effects of oxytocin and TGOT in the EPN, we crossed the OXTR-EGFP transgene onto an *Oxtr*^*-/-*^ line and probed the function of the EPN in response to TGOT (n=10, 6 animals-4 male/2 female). In the absence of functional OXTRs, there was no significant main effect of TGOT bath application on the RMP (F (1.793, 16.13) = 2.628, p=0.107, Supplementary Table 1 Line F), membrane resistance (F (1.429, 12.86) = 0.194, p=0.753, Supplementary Table 1 Line G) or the number of evoked action potentials (F (1.552, 13.97) = 0.778, p=0.447, Table 1 Supplementary Line H) (Figure 4K-P). These data suggest that TGOT and OXT have a depolarizing effect on EPN neurons in an OXTR dependent manner. However, these data do not show whether this is a direct effect of TGOT/OXT or an indirect effect resulting from the excitation or inhibition of other cells within the tissue slice.

Further experiments were performed under glutamate and GABA receptor blockade to investigate whether the OXTR effects are direct or indirect (n=13, 5 animals-3 male/2 female). Initially, an aCSF solution containing picrotoxin (1 μM) and a combination of DNQX (5 μM) and APV (10 μM) was used to block GABA and glutamate receptors in the tissue slice before, during and after TGOT bath application. Under these conditions, TGOT had a similar effect on the RMP, membrane resistance, and number of evoked action potentials seen previously with only TGOT. Specifically, TGOT bath application had a significant main effect on RMP (F (2.070, 24.84) = 6.388, p=0.005) with significantly higher RMP at 4 (Dunnett MCT, MΔ= 3.379 mV, SE=1.258, Q=2.686, df=12, p=0.0497, d=0.444) and 6 minutes (Dunnett MCT, MΔ=4.65, SE=1.466, Q=3.172, df=12, p=0.021, d=0.606) after the start of TGOT application with no significant difference between baseline and washout RMP (Dunnett MCT, MΔ=0.089, SE=1.433, Q=0.062, df=12, p=0.999)(Figure 5A, D-G, Supplementary Table 1 Line I). A main effect of TGOT on membrane resistance was also identified (F (1.583, 19.00) = 5.443, p=0.019) with a significantly higher membrane resistance at 4 (Dunnett MCT, MΔ=45.32 mΩ, SE=16.37, Q=2.768, df=12, p=0.043, d=0.243) and 6 (Dunnett MCT, MΔ=66.34 mΩ, SE=17.45, Q=3.801, df=12, p=0.007, d=0.347) minutes into to TGOT application with a return to baseline values upon washout (Dunnett MCT, MΔ=3.091 mΩ, SE=19.07, Q=0.162, df=12, p=0.997)(Figure 5B, Supplementary Table 1 Line J). Lastly, TGOT also had a significant main effect on the number of evoked action potentials under GABA and glutamate receptor blockade (F (1.610, 19.32) = 4.868, P=0.025). Specifically, the number of evoked action potentials was higher at both 4 (Dunnett MCT, MΔ=3.846, SE=0.815, Q=4.718, df=12, p=0.001, d=0.53) and 6 minutes relative to baseline (Dunnett MCT, MΔ=6.154, SE=1.213, Q=5.071, df=12, p=0.001) with no significant difference between baseline and washout (Dunnett MCT, MΔ=1.154, SE=2.364, Q=0.488, df=12, p=0.928, d=0.753) (Figure 5C-G, Supplementary Table 1 Line K). These data show that the effect of TGOT on EPN neurons is not dependent on activation of either GABA or glutamate receptors via local connections from other neurons, suggesting a more direct effect of TGOT on the EPN neurons. However, OXTR has been found on neurons expressing other neurotransmitters, including serotonergic neurons of the raphe nucleus ^44^ and somatostatin containing neurons in the medial prefrontal, visual and auditory cortices ^11,45,46^, therefore, other neurotransmitters could be mediating the effect of TGOT and OXT application in these experiments.

**Figure 5.**
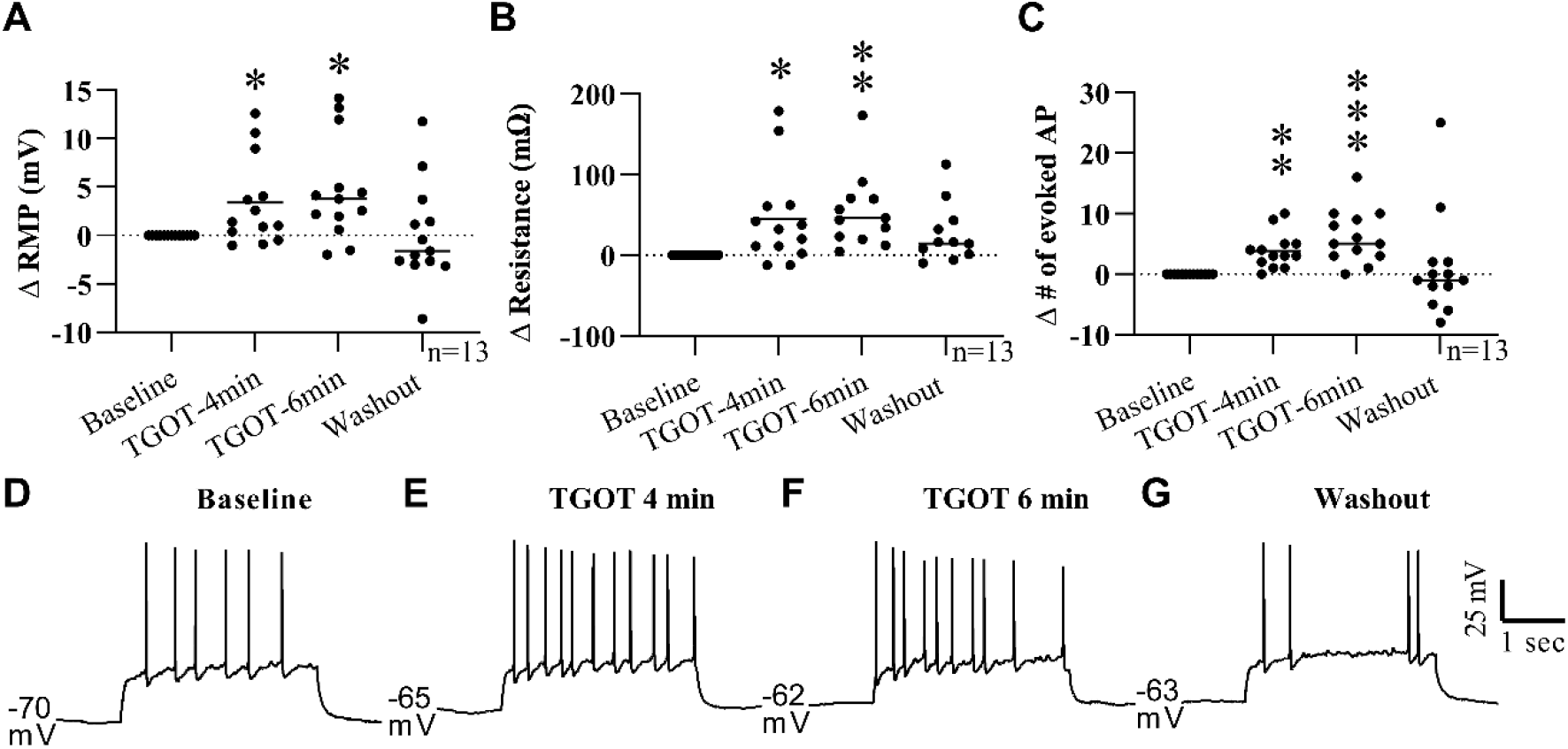
TGOT induced depolarization of EPN neurons persists under GABA and glutamate receptor blockade. To limit indirect activity, GABA and glutamate receptors in the tissue slice were blocked during the baseline recording using bath application of picrotoxin (200 nM), DNQX (5 μM) and APV (10 μM) respectively. The addition of TGOT (200 nM) significantly depolarized the membrane potential (A), increased the membrane resistance (B) and the number of evoked action potentials at 4 and 6 minutes after the start of bath application with a return to baseline levels upon washout in all parameters. Bath application of picrotoxin, DNQX and APV were continued throughout the entire recording. Traces from a representative single EPN cell are shown in D-G. * p<0.05; ** p<0.005; *** p<0.001.

To rule out indirect effects from other neurotransmitters, further experiments using tetrodotoxin to block all synaptic activity were conducted to test the direct effect of TGOT on EPN OXTR-expressing neurons (n=19, 8 animals-3 male/5 female). TTX was added to the aCSF and was started ∼4 minutes prior to the addition of TGOT. Bath application of TTX was continued throughout the recording (i.e. baseline, TGOT and washout). In the absence of synaptic activity, the depolarizing effect of TGOT persisted, indicating a direct effect of TGOT on the recorded EPN neuron. A main effect of TGOT bath application on RMP (F (1.795, 32.31) = 19.50, p<0.0001, RM one-way ANOVA) and membrane resistance (F (1.507, 27.12) = 6.342, p=0.0095) was found using RM one-way ANOVAs. Dunnett MCT post hoc tests indicated a significant increase in the RMP at both 4 (MΔ= 1.957 mV, SE= 0.518, Q=3.778, df=18, p=0.004, d=0.404) and 6 minutes (MΔ=2.007 mV, SE=0.393, Q=5.103, df=18, p=0.0002, d=0.407) after the start of TGOT bath application and a significant hyperpolarization of the membrane potential upon washout (MΔ= -2.343 mV, SE=0.782, Q=3.012, df=18, p=0.02, d=0.409)(Figure 6A,C, Supplementary Table 1 Line L). A significant increase in membrane resistance above baseline levels was also found (Dunnett MCT) at 4 (MΔ=26.9 mΩ, SE=8.746, Q=3.075, df=18, p=0.017, d=0.166) and 6 minutes (MΔ=32.99 mΩ, SE=10.65, Q=3.098, df=18, p=0.017, d=0.202) post TGOT application with no significant difference between baseline and washout levels (MΔ=-23.07 mΩ, SE=17.87, Q=1.291, df=18, p=0.447) (Figure 6B, D, Supplementary Table 1 Line M).

**Figure 6.**
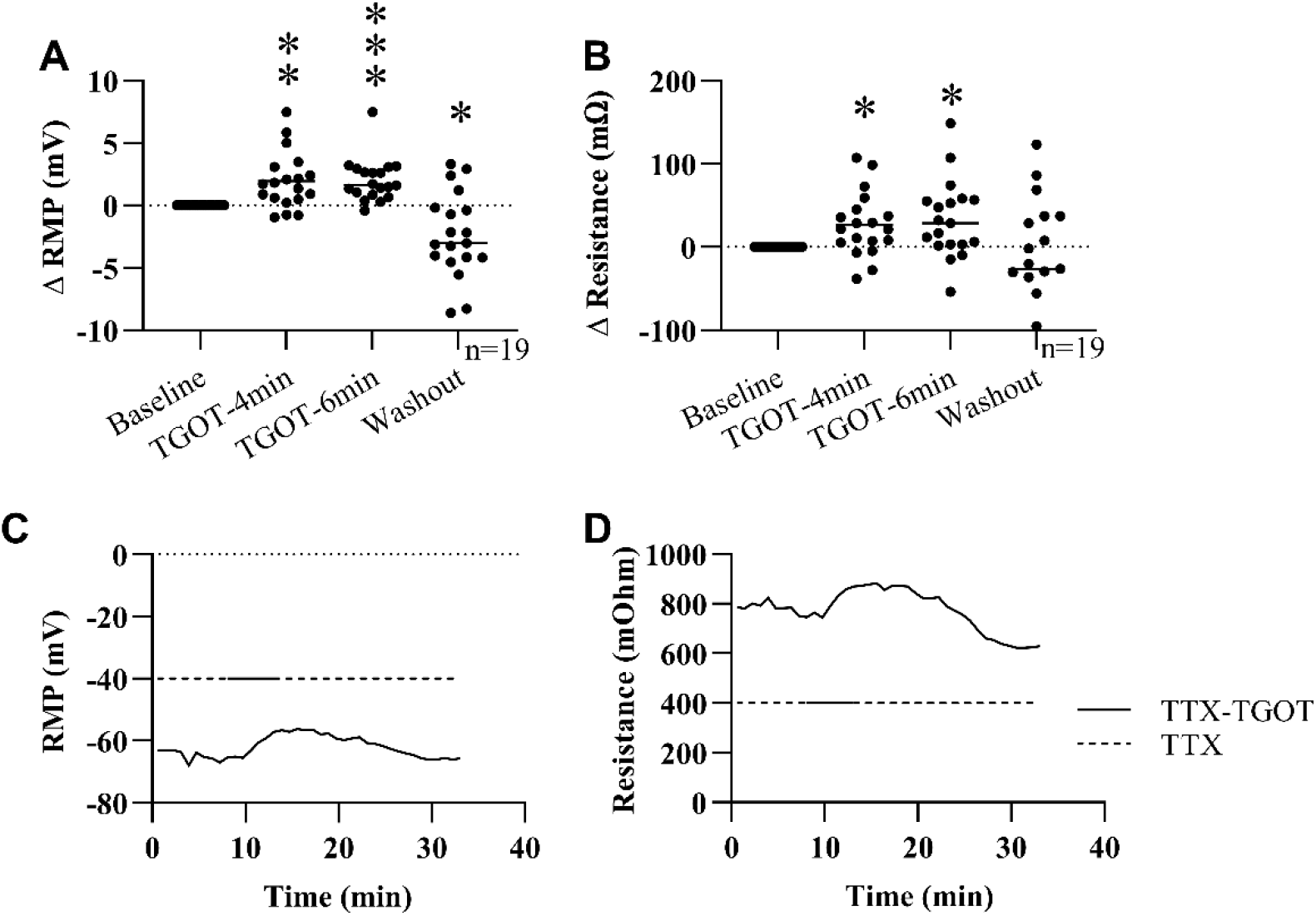
Depolarizing effect of TGOT on EPN neurons persists in the absence of network activity. The direct effect of TGOT on EPN neurons was investigated by using tetrodotoxin (TTX, 1μM) to block all sodium channel activity. In the absence of network activity, bath application of TGOT depolarized the membrane potential (A) and increased the cells membrane resistance (B) at 4 and 6 minutes after the start of bath application. A significant hyperpolarization of the membrane potential was detected under these conditions as well (A). Representative data from a single EPN neuron shows membrane potential (C) and resistance (D) values collected once every sweep (50 seconds) during TTX/TGOT protocol. * p<0.05; ** p<0.005; *** p<0.001.

## Discussion

The experiments outlined in this paper were designed to test the hypothesis that OXTRs are expressed on a specific subpopulation of neurons in the mouse EPN and that OXT can modulate the activity of these OXTR-expressing neurons of the EPN. Using RNAscope in situ hybridization, we found a high level of co-localization of *Oxtr* mRNA and VGlut1 markers in EPN, suggesting that OXTR are highly expressed on projection neurons within the EPN. To test the effect of OXT on EPN neurons, we used whole-cell patch clamp electrophysiology in a genetic mouse model that expresses EGFP in OXTR-expressing neurons. EPN neurons displayed a depolarizing effect of OXT and the OXTR-specific ligand, TGOT in P21-P28 mice. To ensure the effect of TGOT on EPN neurons was OXTR dependent, we also tested the effect of TGOT on *Oxtr* knock-out EPN tissue and found that TGOT did not have a significant effect on EPN cells from *Oxtr* knock-out tissue slices.

In the experiments detailed here, a high co-localization of *Oxtr* and VGLUT1 was found in the EPN neurons, with sparse co-localization of *Oxtr+* and *Gad1* and *Gad2* mRNA. The molecular phenotype of the OXTR-expressing cells determines the indirect effect of OXT on downstream neural targets. For example, in the hippocampus and spinal cord, OXT activation of presynaptic OXTRs facilitates glutamate release from presumably OXTR-expressing glutamatergic neurons ^47,48^ thus having an excitatory effect on both the OXTR expressing cell and its downstream target. In other studies, OXT has also been found to modulate cells expressing the inhibitory neurotransmitter, GABA, in several brain regions, including neocortex, piriform, auditory cortex, hippocampus, hypothalamus and olfactory bulb ^11,12,49–52^. In these examples, OXT can have an inhibitory effect on neurons that receive input from OXTR-expressing, GABAergic cells and could modulate the local excitatory:inhibitory balance. In the auditory cortex, OXT enhances a dam’s pup retrieval in response to calls from a distressed pup. OXT injection into the auditory cortex of virgin dams enhances the auditory signal by balancing inhibitory and excitatory activity which made the spike activity more consistent during the call and strengthened the response resulting in behavior similar to an experienced dam ^11^. Similar fine-tuning of action potential firing by OXT has been documented in other brain areas, including hippocampus ^52^, the main olfactory bulb ^5^ and visual cortex ^45^.

Overall, OXT and the OXTR specific agonist TGOT were found to have a depolarizing effect on EPN neurons. In normal aCSF, the addition of OXT increased the spontaneous firing rate of EPN neurons artificially held near threshold. Since OXT is structurally similar to another neuropeptide, vasopressin, and is able to bind to vasopressin receptors ^39–43^, the OXTR specific agonist, TGOT was used to avoid any confounding effect of vasopressin receptor activation by OXT. TGOT application had a similar effect on EPN neurons in aCSF, again increasing the number of spontaneous action potentials when neurons were held near threshold. In subsequent experiments aimed at further understanding the effect of OXTR activation, TGOT application increased the resting membrane potential, the number of evoked action potentials and increased the membrane resistance. Under these conditions, it is possible that the effects seen were due to OXT action on neurons presynaptic to the recorded neuron, so glutamate and GABA receptor antagonists were used to block any OXT-mediated increase in pre-synaptic glutamate and GABA release. Similar to aCSF, bath application of TGOT with GABA and glutamate antagonists exhibited increased neuronal excitability on EPN neurons. Other neurotransmitters, specifically somatostatin, are known to co-localize with OXTR in other brain areas, e.g. primary visual cortex ^45^. Based on the possibility of OXT modulating other neurotransmitters presynaptically, a final set of experiments were conducted using TTX to block all network activity. Again, under network blockade TGOT increased the excitability of EPN neurons as indicated by a depolarized membrane potential concurrent with an increase in the membrane resistance. In other brain areas, OXT has been shown to have a depolarizing effect using several different GPCR-mediated pathways. In cultured rat supraoptic neurons, OXTRs work via activation of the G_*α*q_ pathway which activates phospholipase C in order to stimulate intracellular calcium release ^53^. Alternatively, activation of the G_q/11_ subpopulation of OXTRs is associated with an inhibition of inward rectifying potassium channels which lead to membrane depolarization and a decrease in conductance of potassium across the membrane ^51,54–56^. In this study, OXT and TGOT bath application resulted in a significant increase in the membrane resistance coupled with a significant membrane depolarization in mouse EPN neurons. These results suggest that OXTRs in EPN may be modulating activity via inhibition of a membrane bound channel, possibly through an inward rectifying potassium channel as seen in previous experiments ^51,55,56^. Further experiments are necessary to examine whether this is the mechanism by which OXT is affecting membrane excitability in the EPN.

Another experiment was performed here to confirm the depolarizing effect of TGOT on EPN neurons. To retain the ability to use the EGFP reporter to identify EPN neurons in acute slices, the OXTR-EGFP mouse line was crossed with an OXTR-KO line to get tissue slices in which there are no functional OXTRs, but cells that would normally express these receptors in a wild-type animal express the EGFP marker. In these animals, the lack of OXTRs should eliminate any effect of TGOT in EGFP-expressing EPN neurons if the effect is mediated by OXTR activation. As expected, TGOT application had no effect on any of the parameters measured (Figure 4) in OXTR knockout slices, confirming the OXT-mediated effect on OXTR-EGFP expressing neurons in EPN was in due to activation of the OXTR.

Although little is known about the role of oxytocin in EPN, it may play an important role in the development of social behavior. Interestingly, OXT has been shown to contribute to both long term potentiation and long term depression of neuronal activity. In several areas of the brain, including the lateral amygdala ^57^, dentate gyrus ^58^, and dorsal horn nociceptive neurons ^59^, OXT elicits long-term depression. However, in other brain areas like the accessory olfactory bulb and hippocampus, OXT facilitates learning and memory ^60^. OXT also enhances spatial memory in mouse dams ^24^ and can block and rescue stress induced deficits in synaptic plasticity ^61,62^. In mice, early maternal separation for three hours a day (PND 1-21) produces social behavior impairments and deficits in learning and memory and the production of hippocampal LTP, which can be rescued by giving intranasal OXT for 10 days following maternal separation (PND 22-34) ^63^. Oxytocin receptors in the hippocampus are also necessary for long-term social recognition in mice as deletion of CA2/CA3 OXTR impairs long-term social-recognition memory and blocks the induction of hippocampal LTP ^22^. Based on the role of OXT in LTP and learning and memory in other brain areas, it is possible that OXTRs in EPN are contributing to learning and memory of social information within EPN.

The EPN receives input from several areas involved in processing the emotional salience of incoming sensory information, including amygdalar areas (anterior cortical amygdala, periamygdaloid complex and medial amygdala) and the infralimbic cortex ^64,65^. EPN also receives input from olfactory and gustatory cortices and this information is integrated within the EPN ^66,67^. EPN also has reciprocal connections with areas involved with learning and memory, specifically the entorhinal cortex which is involved with spatial memory and the perirhinal cortex which receives input regarding sensory information ^64^. Given the reciprocal connections with the areas that process incoming sensory information, emotional salience and memory, EPN is a prime candidate to act as an association area, playing a role in emotional learning of chemosensory information ^64,66^. The input of OXT via the abundant and stable OXTR expression through development may permit the OXT system to facilitate learning and memory of emotionally salient sensory stimuli during development, thus modifying the development of adult social behaviors. For example, in the presence of OXT, incoming sensory information from the olfactory system may be more likely to induce long-term potentiation and memory formation within the EPN neurons because of the depolarizing effect of OXT.

The OXTR-mediated depolarizing effect of OXT in EPN neurons has been demonstrated here, however because little is known about the function of the EPN it is still unclear how this OXTR mediated effect can modulate behavior. It is possible that the EPN is important for synaptic plasticity and learning of social behaviors throughout development. For example, in situations where OXT is released, i.e. suckling, dam-infant caregiving, social interactions in juvenile and adult mice, OXT action in the EPN may be modulating synaptic connections or downstream excitatory/inhibitory balance to elicit long-term changes in the circuitry. The data here show that OXTR activation from P21-P28 is excitatory to presumably mainly glutamatergic neurons, thus it likely affects downstream brain areas. Evidence suggests that EPN is important for integrating sensory information from the olfactory and gustatory system ^66,67^ and with the data presented here, it may also be integrating OXT signaling which could modulate the salience of incoming sensory information and pass this information on to downstream limbic areas like the amygdala in order to modulate social behavior. Additional experiments are necessary to determine the full extent of OXT-mediated activity in EPN and its downstream neuronal targets, and impact on behavior.

## Acknowledgements

The authors would like to acknowledge Kristen Abreu, Esther Rodriguez-Forti, and Natalia Fernandez, as well as the animal care staff at Florida State University.

## Author contributions

LMB and EADH designed research and wrote the paper. LMB also performed research and analyzed data.

## Data availability

All data generated or analyzed during this study are included in this published article, its Supplementary Information files and in the GenBank genetic sequence database (NM_001081147.1, NM_008077.4, NM_008078.2, NM_182993.2, NM_080853.3).

