## Supplementals Tables 1-14 for "Oxytocin via oxytocin receptor excites neurons in the endopiriform nucleus of juvenile mice"

### Supplementary material for "Oxytocin via oxytocin receptor excites neurons in the endopiriform nucleus of juvenile mice"

Authors: Lindsey M. Biggs and Elizabeth A.D. Hammock

**Supplementary Table 1: Statistical analyses.** Table shows the statistical analyses used for each figure in the main text. Data was analyzed at the level of the individual cell. In some cases, data from more than one cell was collected per animal so the number and sex of animals included in the sample size is included in the table. Once an experimental bath was applied to the slice, it was discarded so all cells were naïve to the experimental drug baths at the start of the recording.

| | Figure # | Data Structure | Statistical test | Sample Size | Statistical Results | Post-hoc test results |
|---|---|---|---|---|---|---|
| A | 4A | Normal Distribution | One-way RM ANOVA with Geisser-Greenhous correction (GG) | n = 6 cells, 3 animals: 1 male, 2 female | F = 6.315, p = 0.015 | Dunnett MCT: Baseline vs OXT 4 min: p = 0.011, d = 1.87; Baseline vs OXT 6 min: p = 0.253; Baseline vs Washout: p = 0.768 |
| B | 4B | Normal Distribution | One-way RM ANOVA with GG correction | n = 8 cells, 2 animals: 1 male, 1 female | F = 8.918, p = 0.008 | Dunnett MCT: Baseline vs OXT 4 min: p = 0.036, d = 1.25; Baseline vs OXT 6 min: p = 0.246; Baseline vs Washout: p = 0.014, d = 1.35 |
| C | 4D | Normal Distribution | One-way RM ANOVA with GG correction | n = 11 cells, 4 animals: 2 male, 2 female | F = 6.165, p = 0.006 | Dunnett MCT: Baseline vs OXT 4 min: p = 0.019, d = 0.275; Baseline vs OXT 6 min: p = 0.022, d = 0.379; Baseline vs Washout: p = 0.72 |
| D | 4E | Normal Distribution | One-way RM ANOVA with GG correction | n = 11 cells, 4 animals: 2 male, 2 female | F = 5.533, p = 0.014 | Dunnett MCT: Baseline vs OXT 4 min: p = 0.395; Baseline vs OXT 6 min: p = 0.001, d = 0.518; Baseline vs Washout: p = 0.682 |
| E | 4F | Normal Distribution | One-way RM ANOVA with GG correction | n = 11 cells, 4 animals: 2 male, 2 female | F = 5.426, p = 0.018 | Dunnett MCT: Baseline vs OXT 4 min: p = 0.003, d = 1.185; Baseline vs OXT 6 min: p = .009, d = 1.739; Baseline vs Washout: p > 0.999 |
| F | 4K | Normal Distribution | One-way RM ANOVA with GG correction | n = 10 cells, 6 animals: 4 male, 2 female | F = 2.628, p = 0.107 | NA |
| G | 4L | Normal Distribution | One-way RM ANOVA | n = 10 cells, 6 animals: | F = 0.194, p = 0.753 | NA |

| | | | | | | |
|---|---|---|---|---|---|---|
| | | | with GG correction | 4 male, 2 female | | |
| H | 4M | Normal Distribution | One-way RM ANOVA with GG correction | n = 10 cells, 6 animals: 4 male, 2 female | F = 0.778, p = 0.447 | NA |
| I | 5A | Normal Distribution | One-way RM ANOVA with GG correction | n = 13 cells, 5 animals: 3 male, 2 female | F = 6.388, p = 0.005 | Dunnett MCT: Baseline vs OXT 4 min: p = 0.0497, d = 0.444; Baseline vs OXT 6 min: p = 0.021, d = 0.606, Baseline vs Washout: p = 0.999 |
| J | 5B | Normal Distribution | One-way RM ANOVA with GG correction | n = 13 cells, 5 animals: 3 male, 2 female | F = 5.443, p = 0.019 | Dunnett MCT: Baseline vs OXT 4 min: p = 0.043, d = 0.243; Baseline vs OXT 6 min: p = 0.007, d = 0.347; Baseline vs Washout: p = 0.997 |
| K | 5C | Normal Distribution | One-way RM ANOVA with GG correction | n = 13 cells, 5 animals: 3 male, 2 female | F = 4.868, p = 0.025 | Dunnett MCT: Baseline vs OXT 4 min: p = 0.001, d = 0.53; Baseline vs OXT 6 min: p = 0.001, d = 0.753; Baseline vs Washout: p = 0.928 |
| L | 6A | Normal Distribution | One-way RM ANOVA with GG correction | n = 19 cells, 8 animals: 3 male, 5 female | F = 19.5, p < 0.0001 | Dunnett MCT: Baseline vs OXT 4 min: p = 0.004, d = 0.404; Baseline vs OXT 6 min: p = 0.0002, d = 0.407; Baseline vs Washout: p = 0.02, d = 0.409 |
| M | 6B | Normal Distribution | One-way RM ANOVA with GG correction | n = 19 cells, 8 animals: 3 male, 5 female | F = 6.342, p = 0.01 | Dunnett MCT: Baseline vs OXT 4 min: p = 0.017, d = 0.166; Baseline vs OXT 6 min: p = 0.017, d = 0.202; Baseline vs Washout: p = 0.447 |

**Supplementary Table 2: Oxytocin effect on the number of spontaneous action potentials (APs) in cells held near threshold.** Data presented as the absolute change from baseline value. For each cell, the baseline number of spontaneous action potentials was subtracted from the number of spontaneous action potentials during the experimental bath indicated. Data used for analysis of Figure 4A.

| Cell ID | Baseline (normalized to 0) | 4 min: Δ# of AP (4min-baseline) | 6 min: Δ# of AP (6min-baseline) | Washout (30 min): Δ# of AP (Washout-baseline) |
|---|---|---|---|---|
| 03182019-2-1 | 0 | 30 | 49 | -9 |
| 03212019-1-1 | 0 | 57 | 69 | 5 |
| 03252019-2-1 | 0 | 42 | -19 | -28 |
| 03182019-2-3 | 0 | 27 | 36 | 24 |
| 03212019-1-2 | 0 | 7 | -3 | -3 |
| 03252019-2-2 | 0 | 34 | 19 | -29 |

**Supplementary Table 3: TGOT effect on the number of spontaneous APs in cells held near threshold.** Data presented as the absolute change from baseline value. For each cell, the baseline number of spontaneous APs was subtracted from the number of spontaneous APs during the experimental bath indicated. Data used for analysis of Figure 4B.

| Cell ID | Baseline (normalized to 0) | 4 min: Δ# of AP (4min-baseline) | 6 min: Δ# of AP (6min-baseline) | Washout: Δ# of AP (Washout-baseline) |
|---|---|---|---|---|
| 04242019-1-1 | 0 | -3 | -3.5 | -4.5 |
| 04242019-1-2 | 0 | 21 | 13 | -1.5 |
| 04242019-1-3 | 0 | 37 | 23.5 | -11.5 |
| 10042019-2-1 | 0 | 15.5 | 20 | -6.5 |
| 04242019-1-5 | 0 | 29 | 6.5 | -3.5 |
| 04242019-1-6 | 0 | 43.5 | 64.5 | -7 |
| 10042019-2-2 | 0 | 24.5 | 8.5 | -18 |
| 10042019-2-3 | 0 | -6.5 | -13 | -17.5 |

**Supplementary Table 4: TGOT effect on the resting membrane potential (RMP) of cells with no holding current applied.** Data presented as the absolute change from baseline value. For each cell, the baseline RMP was subtracted from the RMP during the experimental bath indicated. Data used for analysis of Figure 4D.

| **Cell ID** | **Baseline RMP (normalized to 0)** | **4 min: ΔRMP (mV) (4min-Baseline)** | **6 min: ΔRMP (mV) (6min-Baseline)** | **Washout: ΔRMP (Washout -Baseline)** |
|---|---|---|---|---|
| 05282019-1-3 | 0 | 0.696014404 | 2.616638184 | 1.485603333 |
| 05292019-2-3 | 0 | 2.827163696 | 0.539176941 | 1.483146667 |
| 05292019-2-4 | 0 | 1.646781921 | 6.568389893 | 0.46572113 |
| 05312019-1-1 | 0 | 1.642494202 | 2.380065918 | 5.337211609 |
| 05282019-1-1 | 0 | 2.591697693 | 1.735923767 | -3.197929382 |
| 05282019-1-4 | 0 | 1.636474609 | 3.605308533 | 1.578430176 |
| 05292019-2-1 | 0 | 4.82686615 | 10.16745758 | 1.4323349 |
| 05292019-2-2 | 0 | -0.624221802 | 1.161521912 | -2.684371948 |
| 05302019-1-3 | 0 | 0.140625 | 0.165588379 | -2.010498047 |
| 05312019-1-3 | 0 | 7.757534027 | 7.926109314 | 1.602947235 |
| 05312019-1-4 | 0 | 2.710834503 | 0.039001465 | 1.597640991 |

**Supplementary Table 5: TGOT effect on the membrane resistance (mΩ) of cells with no holding current applied.** Data presented as the absolute change from baseline value. For each cell, the baseline membrane resistance was subtracted from the membrane resistance during the experimental bath indicated. Data used for analysis of Figure 4E.

| **Cell ID** | **Baseline Resistance (mΩ) (normalized to 0)** | **4 min: ΔResistance (mΩ) (4min-Baseline)** | **6 min: ΔResistance (mΩ) (6min-Baseline)** | **Washout: ΔResistance (mΩ) (Washout-Baseline)** |
|---|---|---|---|---|
| 05282019-1-3 | 0 | 14.58374023 | 52.74810791 | 82.5579834 |
| 05292019-2-3 | 0 | 10.28945923 | 67.52059937 | 7.488098145 |
| 05292019-2-4 | 0 | 36.64117432 | 160.4873047 | -27.64929199 |
| 05312019-1-1 | 0 | 35.64562988 | 59.28649902 | 89.65576172 |
| 05282019-1-1 | 0 | -29.7315979 | 43.52920532 | 87.12939453 |
| 05282019-1-4 | 0 | 17.10968018 | 14.79064941 | -51.74697876 |
| 05292019-2-1 | 0 | 24.2539978 | 96.58425903 | 18.02429199 |
| 05292019-2-2 | 0 | 15.22015381 | 68.0960083 | -2.096740723 |
| 05302019-1-3 | 0 | 10.78018188 | 20.47210693 | -9.259796143 |
| 05312019-1-3 | 0 | 125.8762512 | 92.5930481 | -33.42858887 |
| 05312019-1-4 | 0 | -52.49676514 | 38.06970215 | -4.746551514 |

**Supplementary Table 6: TGOT effect on the number of APs evoked by a two second intracellular current injection.** Value of current injection was determined for each cell prior to the start of data collection such that multiple APs were evoked (between 2-10 evoked APs) to avoid floor and ceiling effects. Data presented as the absolute change from baseline value. For each cell, the baseline number of evoked APs was subtracted from the number of APs during the experimental bath indicated. Data used for analysis of Figure 4F.

| **Cell ID** | **Baseline # of evoked AP (normalized to 0)** | **4 min: Δ # of evoked AP (4min-Baseline)** | **6 min: Δ # of evoked AP (6min-Baseline)** | **Washout: Δ # of evoked AP [Washout -Baseline]** |
|---|---|---|---|---|
| 05282019-1-3 | 0 | 0 | 1 | 3 |
| 05292019-2-3 | 0 | 5 | 6 | -2 |
| 05292019-2-4 | 0 | 1 | -1 | 1 |
| 05312019-1-1 | 0 | 3 | 5 | 6 |
| 05282019-1-1 | 0 | 3 | 0 | -1 |
| 05282019-1-4 | 0 | 1 | 6 | 1 |
| 05292019-2-1 | 0 | 6 | 5 | -3 |
| 05292019-2-2 | 0 | 2 | 6 | 7 |
| 05302019-1-3 | 0 | 7 | 12 | 4 |
| 05312019-1-3 | 0 | 6 | 6 | -6 |
| 05312019-1-4 | 0 | 2 | 1 | -10 |

**Supplementary Table 7: TGOT effect on the RMP (mV) of cells from OXTR-KO tissue with no holding current applied.** Data presented as the absolute change from baseline value. For each cell, the baseline RMP was subtracted from the RMP during the experimental bath indicated. Data used for analysis of Figure 4K.

| **Cell ID** | **Baseline RMP (mV) (normalized to 0)** | **4 min: ΔRMP (mV) (4min-Baseline)** | **6 min: ΔRMP (mV) (6min-Baseline)** |
|---|---|---|---|
| 12232019-1-1 | 0 | -0.693328857 | -1.93396759 |
| 12232019-1-3 | 0 | -2.166801453 | -0.376533508 |
| 12232019-1-4 | 0 | 0.391983032 | -0.35471344 |
| 01072020-1-3 | 0 | -7.566902161 | -7.031013489 |
| 01082020-1-2 | 0 | -3.481842041 | 1.260932922 |
| 01082020-1-3 | 0 | -6.31060791 | -4.068275452 |
| 01102020-1-1 | 0 | 0.793952942 | -2.843727112 |
| 01102020-1-2 | 0 | 1.096287964 | 0.381650208 |
| 01132020-1-2 | 0 | -0.200508118 | 0.748931885 |
| 02042020-1-1 | 0 | -0.159294128 | 0.038200378 |

**Supplementary Table 8: TGOT effect on the membrane resistance (mΩ) of OXTR-KO cells with no holding current applied.** Data presented as the absolute change from baseline value. For each cell, the baseline membrane resistance was subtracted from the membrane resistance during the experimental bath indicated. Data used for analysis of Figure 4L.

| Cell ID | Baseline Resistance (mΩ) (normalized to 0) | 4 min: ΔResistance (mΩ) (4min-Baseline) | 6 min: ΔResistance (mΩ) (6min-Baseline) |
|---|---|---|---|
| 12232019-1-1 | 0 | 3.869628906 | -10.13870239 |
| 12232019-1-3 | 0 | -71.50524902 | -12.82406616 |
| 12232019-1-4 | 0 | 14.22576904 | 0.046234131 |
| 01072020-1-3 | 0 | -62.29217529 | -74.67697144 |
| 01082020-1-2 | 0 | 13.78967285 | 22.7479248 |
| 01082020-1-3 | 0 | 26.79199219 | -0.336303711 |
| 01102020-1-1 | 0 | 71.8230896 | 49.43664551 |
| 01102020-1-2 | 0 | -76.90200806 | -34.25872803 |
| 01132020-1-2 | 0 | 4.048309326 | 4.122924805 |
| 02042020-1-1 | 0 | 2.083282471 | 21.20849609 |

**Supplementary Table 9: TGOT effect on the number of APs evoked by a two second intracellular current injection in OXTR-KO cells.** Value of current injection was determined for each cell prior to the start of data collection such that multiple APs were evoked (between 2-12 evoked APs) to avoid floor and ceiling effects. Data presented as the absolute change from baseline value. For each cell, the baseline number of evoked APs was subtracted from the number of APs during the experimental bath indicated. Data used for analysis of Figure 4M.

| Cell ID | Baseline # of evoked AP (normalized to 0) | 4 min: Δ # of evoked AP (4min-Baseline) | 6 min: Δ # of evoked AP (6min-Baseline) |
|---|---|---|---|
| 12232019-1-1 | 0 | 1 | 2 |
| 12232019-1-3 | 0 | 1 | 0 |
| 12232019-1-4 | 0 | 1 | 2 |
| 01072020-1-3 | 0 | -3 | -2 |
| 01082020-1-2 | 0 | 0 | 0 |
| 01082020-1-3 | 0 | -5 | -4 |
| 01102020-1-1 | 0 | -1 | -3 |
| 01102020-1-2 | 0 | 0 | 0 |
| 01132020-1-2 | 0 | 0 | 2 |
| 02042020-1-1 | 0 | -1 | 1 |

**Supplementary Table 10: TGOT effect on the RMP of cells under glutamate and GABA receptor blockade (DNQX:5 µM, APV: 10 µM, picrotoxin:200 nM) with no holding current applied.** Data presented as the absolute change from baseline value (mV). For each cell, the baseline RMP was subtracted from the RMP during the experimental bath indicated. Data used for analysis of Figure 5A.

| Cell ID | Baseline RMP (mV) (normalized to 0) | 4 min: ΔRMP (mV) (4min-Baseline) | 6 min: ΔRMP (mV) (6min-Baseline) | Washout: ΔRMP (mV) (Washout-Baseline) |
|---|---|---|---|---|
| 05132019-1-4 | 0 | 2.614860535 | 4.441215515 | -3.023742676 |
| 05142019-1-2 | 0 | 1.037597656 | 2.560382843 | -0.432037354 |
| 05092019-1-2 | 0 | 10.62329102 | 13.20077133 | 7.15184021 |
| 05102019-1-1 | 0 | 8.972099304 | 14.1973381 | 11.76184464 |
| 05102019-1-2 | 0 | 0.878650665 | 1.97864151 | -8.608024597 |
| 05102019-1-3 | 0 | 12.60706711 | 11.95859528 | -2.652645111 |
| 05152019-1-2 | 0 | -0.470970154 | 2.177902222 | -1.611412048 |
| 05152019-1-3 | 0 | 1.394363403 | 4.136558533 | -2.614006042 |
| 05132019-1-2 | 0 | 0.414199829 | 0.590335846 | -2.006576538 |
| 05142019-1-3 | 0 | 3.685409546 | 4.94493866 | 1.160240173 |
| 05142019-1-4 | 0 | -1.000091553 | -1.991653442 | 3.716644287 |
| 05102019-1-4 | 0 | 4.070259094 | 3.790687561 | -3.143341064 |
| 05152019-1-4 | 0 | -0.903282166 | -1.537826538 | 1.455093384 |

**Supplementary Table 11: TGOT effect on the membrane resistance (mΩ) of cells under glutamate and GABA receptor blockade (DNQX:5 µM, APV: 10 µM, picrotoxin:200 nM) with no holding current applied.** Data presented as the absolute change from baseline value. For each cell, the baseline membrane resistance was subtracted from the membrane resistance during the experimental bath indicated. Data used for analysis of Figure 5B.

| Cell ID | Baseline Resistance (mΩ) (normalized to 0) | 4 min: ΔResistance (mΩ) (4min-Baseline) | 6 min: ΔResistance (mΩ) (6min-Baseline) | Washout: ΔResistance (mΩ) (Washout-Baseline) |
|---|---|---|---|---|
| 05132019-1-4 | 0 | 32.47894287 | 70.64300537 | -124.1886749 |
| 05142019-1-2 | 0 | 20.34307861 | 19.85501099 | 14.40634155 |
| 05092019-1-2 | 0 | 154.1160889 | 216.7189941 | 73.88269043 |
| 05102019-1-1 | 0 | 11.41659546 | 56.56439209 | 43.40658569 |
| 05102019-1-2 | 0 | 62.35211182 | 90.87200928 | -139.1866455 |
| 05102019-1-3 | 0 | 178.5906982 | 173.4503784 | -9.690093994 |
| 05152019-1-2 | 0 | -12.47467041 | 23.2482605 | 16.49533081 |
| 05152019-1-3 | 0 | 42.41680908 | 34.14300537 | -5.99822998 |
| 05132019-1-2 | 0 | 2.28112793 | 4.505859375 | 16.2361145 |
| 05142019-1-3 | 0 | 37.98736572 | 46.48522949 | 32.37243652 |
| 05142019-1-4 | 0 | -12.37243652 | 43.93432617 | 112.7546387 |
| 05102019-1-4 | 0 | 60.81539917 | 69.62713623 | 8.422698975 |
| 05152019-1-4 | 0 | 11.22344971 | 12.32666016 | 1.267486572 |

**Supplementary Table 12: TGOT effect on the number of APs evoked by a two second intracellular current injection under glutamate and GABA receptor blockade (DNQX:5 µM, APV: 10 µM, picrotoxin:200 nM).** Value of current injection was determined for each cell prior to the start of data collection such that multiple APs were evoked (between 2-10 evoked APs) to avoid floor and ceiling effects. Data presented as the absolute change from baseline value. For each cell, the baseline number of evoked APs was subtracted from the number of APs during the experimental bath indicated. Data used for analysis of Figure 5C.

| Cell ID | Baseline # of evoked AP (normalized to 0) | 4 min: Δ # of evoked AP (4min-Baseline) | 6 min: Δ # of evoked AP (6min-Baseline) | Washout: Δ # of evoked AP [Washout-Baseline] |
|---|---|---|---|---|
| 05132019-1-4 | 0 | 2 | 9 | -1 |
| 05142019-1-2 | 0 | 4 | 3 | -8 |
| 05092019-1-2 | 0 | 5 | 4 | -2 |
| 05102019-1-1 | 0 | 1 | 6 | 2 |
| 05102019-1-2 | 0 | 3 | 5 | -6 |
| 05102019-1-3 | 0 | 3 | 1 | -1 |
| 05152019-1-2 | 0 | 4 | 16 | 11 |
| 05152019-1-3 | 0 | 1 | 3 | -5 |
| 05132019-1-2 | 0 | 3 | 10 | 0 |
| 05142019-1-3 | 0 | 10 | 10 | 2 |
| 05142019-1-4 | 0 | 5 | 8 | 25 |
| 05102019-1-4 | 0 | 9 | 5 | 0 |
| 05152019-1-4 | 0 | 0 | 0 | -2 |

**Supplementary Table 13: TGOT effect on the RMP (mV) of cells in the presence of TTX (1 µM) with no holding current applied. Data presented as the absolute change from baseline value.** For each cell, the baseline RMP was subtracted from the RMP during the experimental bath indicated. Data used for analysis of Figure 6A.

| Cell ID | Baseline RMP (mV) (normalized to 0) | 4 min: ΔRMP (mV) (4min-Baseline) | 6 min: ΔRMP (mV) (6min-Baseline) | Washout: ΔRMP (mV) [Washout-Baseline] |
|---|---|---|---|---|
| 06192019-1-2 | 0 | 2.076293945 | 1.349533081 | -3.132415771 |
| 06192019-1-3 | 0 | 7.475727081 | 7.475727081 | -0.714347839 |
| 06202019-1-2 | 0 | 2.156536102 | 1.41431427 | -3.007465363 |
| 06212019-1-3 | 0 | 1.830680847 | 3.22858429 | -0.199333191 |
| 06212019-1-4 | 0 | 0.898830414 | 2.613349915 | 3.324886322 |
| 06242019-1-5 | 0 | 0.910942078 | 0.641773224 | -2.198093414 |
| 06282019-1-3 | 0 | 1.722602844 | 1.731731415 | -4.561096191 |
| 07032019-1-2 | 0 | 5.011901855 | 1.627258301 | -0.368354797 |
| 07022019-1-3 | 0 | 3.097732544 | 3.07056427 | -2.159843445 |
| 07092019-1-1 | 0 | 3.499336243 | 2.659255981 | 2.383552551 |
| 07092019-1-3 | 0 | -0.939495087 | 0.839252472 | 1.221614838 |
| 06282019-1-2 | 0 | 0.496017456 | 3.161933899 | -4.170066833 |
| 07032019-1-3 | 0 | -0.770851135 | -0.426704407 | -4.017044067 |
| 07022019-1-2 | 0 | 5.854957581 | 2.937576294 | -5.558082581 |
| 06202019-1-4 | 0 | 2.425674438 | 0.375320435 | 2.910491943 |
| 06192019-1-1 | 0 | 0.205818176 | 2.624984741 | -8.275993347 |
| 06202019-1-3 | 0 | -0.740913391 | 1.056350708 | -3.241455078 |
| 06212019-1-2 | 0 | 0.606632233 | 1.475036621 | -4.141872406 |
| 06242019-1-4 | 0 | 1.357658386 | 0.27570343 | -8.621170044 |

**Supplementary Table 14: TGOT effect on the membrane resistance (mΩ) of in the presence of TTX (1 μM) with no holding current applied.** Data presented as the absolute change from baseline value. For each cell, the baseline membrane resistance was subtracted from the membrane resistance during the experimental bath indicated. Data used for analysis of Figure 6B.

| Cell ID | Baseline Resistance (mΩ) (normalized to 0) | 4 min: ΔResistance (mΩ) (4min-Baseline) | 6 min: ΔResistance (mΩ) 6min-Baseline) | Washout: ΔResistance (mΩ) (Washout-Baseline) |
|---|---|---|---|---|
| 06192019-1-2 | 0 | -6.655578613 | 1.72076416 | -26.00891113 |
| 06192019-1-3 | 0 | 107.2595215 | 107.2595215 | -122.4525146 |
| 06202019-1-2 | 0 | 98.52502441 | 148.8198242 | 86.13128662 |
| 06212019-1-3 | 0 | 37.29089355 | 55.46832275 | -29.31195068 |
| 06212019-1-4 | 0 | 10.63800049 | 47.73400879 | 68.54412842 |
| 06242019-1-5 | 0 | 72.60980225 | 58.23278809 | -36.38931274 |
| 06282019-1-3 | 0 | 35.83868408 | 16.54803467 | -20.11120605 |
| 07032019-1-2 | 0 | -38.24719238 | 28.96051025 | 123.1519775 |
| 07022019-1-3 | 0 | 45.42984009 | 74.36080933 | -1.886749268 |
| 07092019-1-1 | 0 | 21.24310303 | -9.571533203 | -30.16619873 |
| 07092019-1-3 | 0 | 5.35484314 | 11.75788879 | 7.953643799 |
| 06282019-1-2 | 0 | 21.74603271 | 52.8343811 | 37.30969238 |
| 07032019-1-3 | 0 | -4.851074219 | 6.173553467 | -55.82244873 |
| 07022019-1-2 | 0 | 59.34356689 | 56.72393799 | -95.28070068 |
| 06202019-1-4 | 0 | 28.89868164 | 32.00027466 | 37.44299316 |
| 06192019-1-1 | 0 | -27.77496338 | -53.84738867 | -162.2638245 |
| 06202019-1-3 | 0 | 7.100219727 | 3.007324219 | 28.53436279 |
| 06212019-1-2 | 0 | 29.0201416 | 3.15826416 | -128.7002563 |
| 06242019-1-4 | 0 | 8.290557861 | -14.51782227 | -118.9764099 |
